# Negative regulation of T_H_17-mediated inflammation by the nuclear receptor REV-ERBβ

**DOI:** 10.64898/2026.01.26.701826

**Authors:** Sean Campbell, Sarah A. Mosure, Mohammed Amir, Jonathan Chuck, Qun Lu, Laura A. Solt

## Abstract

T_H_17 cells play a central role in several human autoimmune diseases. We and others reported the nuclear receptor, REV-ERBα, as a cell-intrinsic repressor of T_H_17-mediated pathogenicity. REV-ERBβ, REV-ERBα’s closely related family member, is thought to be functionally redundant to REV-ERBα, which we sought to explore in T_H_17-mediated immunity. Our data indicate that deletion of REV-ERBβ enhances T_H_17-mediated pro-inflammatory cytokine expression and exacerbated disease in mouse models of multiple sclerosis and colitis. RNA-sequencing indicates REV-ERBβ and REV-ERBα do not have similar transcriptional profiles. REV-ERBβ does not appear to regulate gene expression through interaction with the classic co-repressor NCoR1, which is in contrast to REV-ERBα in T_H_17 cells, nor does it utilize heme, its known endogenous ligand for its repressive functions. Our results establish that while REV-ERBβ also acts as a negative regulator of T_H_17-cell function and pathogenicity, it does so in a manner that is non-redundant, independent, and unique to REV-ERBα.

## INTRODUCTION

T_H_17 cells are an IL-17A-secreting subset of CD4^+^ T effector cells (Teff) that defend against invading extracellular pathogens, maintaining immunity at skin and mucosal barriers under homeostatic conditions. In addition to IL-17A, T_H_17 cells are characterized by the production of the cytokines IL-17F, IL-21, and IL-22, eliciting recruitment of other pro-inflammatory mediators to promote pathogen clearance^1^. In contrast, aberrant activation of T_H_17 cells has been associated with the development of several autoimmune and chronic inflammatory diseases, including multiple sclerosis, psoriasis, and inflammatory bowel disease (IBD)^1^. These data suggest that targeting the IL-17 pathway may be a viable therapeutic option for the treatment of these T_H_17-mediated diseases. While IL-17A/IL-17R (receptor) neutralizing antibodies have found success in the treatment of patients with psoriasis, they have had limited success in treating patients with IBD^2^. These data suggest that targeting factors that broadly regulate T_H_17 cell function may be more effective. Therefore, we need a better understanding of the signals and factors that regulate the T_H_17-phenotype to help uncover new therapeutic avenues.

While the nuclear receptor (ΝR) RORγt is considered the lineage defining transcription factor driving T_H_17-cell development, additional data suggest a complex network of transcription factors work in concert to promote or inhibit T_H_17-mediated inflammation^3^. We previously identified that the NR, REV-ERBα, acts as a negative regulator of the T_H_17-mediated program *in vitro* and *in vivo*^4^. Further, REV-ERBα was amenable to ligand-regulation and use of REV-ERB-selective synthetic ligands mediated protection against T_H_17-driven diseases in mouse models of multiple sclerosis (MS) and ulcerative colitis (UC)^4–6^. However, these ligands either targeted both REV-ERBs or REV-ERBα alone. While significantly more is known about REV-ERBα’s physiological function both in and outside the immune system, little is known about the function of REV-ERBβ. In tissues outside of the immune system, both REV-ERBs exhibit similar expression kinetics and target genes, leading many to believe that REV-ERBβ is functionally redundant to REV-ERBα^7–9^. However, in T_H_17 cells, expression of REV-ERBα and REV-ERBβ do not overlap, suggesting that REV-ERBβ may not be functionally redundant to REV-ERBα and have its own unique role^4^. Further, little is known about the role of REV-ERBβ in T cell effector functions, specifically pro-inflammatory T_H_17 cell effector functions and autoimmunity. Therefore, understanding how REV-ERBβ functions will provide insight into the pathophysiology of T_H_17-mediated diseases and determine whether targeting REV-ERBα alone, or in conjunction with REV-ERBβ, is a more advisable option for T_H_17-mediated diseases^4–6^.

In this work, we report that REV-ERBβ is required for dampening T_H_17-mediated pro-inflammatory responses. Genetic deletion of REV-ERBβ resulted in enhanced T_H_17 cell development *in vitro* and exacerbated autoimmune and chronic inflammatory responses *in vivo*. The transcriptional program regulated by REV-ERBβ is largely distinct from REV-ERBα and, in T_H_17 cells, does not appear to utilize the co-repressor NCoR1 (Nuclear Receptor Corepressor 1), a well-defined transcriptional co-factor enabling repressive functions of the REV-ERBs. Collectively, our data suggest that REV-ERBβ functions outside of its classical role as a core member of the circadian clock under pro-inflammatory conditions. Importantly, REV-ERBβ does not appear to be redundant to REV-ERBα functioning as an additional cell-intrinsic negative regulator of T_H_17 cell pro-inflammatory immune responses.

## RESULTS

### Genetic deletion of REV-ERBβ exacerbates pro-inflammatory responses in EAE

To establish whether REV-ERBβ is required to limit T_H_17-mediated pro-inflammatory responses *in vivo*, we used REV-ERBβ^−/−^ mice^10^ to explore the endogenous role of REV-ERBβ in experimental autoimmune encephalomyelitis (EAE), a mouse model of MS. Immune phenotyping revealed no overt differences in thymocyte or T cell composition between REV-ERBβ^+/+^ (wild-type, WT) and REV-ERBβ^−/−^ (knock-out, βKO) mice **(Figure S1A-C**). To induce EAE, we immunized WT and βKO mice with myelin oligodendrocyte glycoprotein peptide (MOG_35-55_) and monitored mice daily for signs of disease. Disease was exacerbated in βKO mice, as demonstrated by a statistically significant difference in Mean Maximum Disease score (score 3.57) relative to WT mice (score 2.05) (**Figure 1A and Table 1**). The mean day of onset of disease symptoms was also earlier in βKO mice compared to WT littermate controls (WT onset, 11.62±1.12; βKO onset, 10.43±1.27) (**Figure 1A and Table 1**). This was likely due to an increase in frequency and number of CD4^+^ T cells infiltrating the central nervous system (CNS) of the βKO mice at the peak of disease (**Figure 1B**). However, the disease phenotype was not attributable to increased frequencies of CD4^+^CCR6^+^ or decreased frequencies of anti-inflammatory Foxp3^+^ T regulatory cells (Tregs), as no difference was observed between WT and βKO mice at peak of disease (Day 15). CCR6 is a chemokine receptor that enables T_H_17 cell migration into inflamed peripheral tissue ^11^ (**Figure 1C, D**). However, there was an increased number of CD4^+^CCR6^+^ T cells in the CNS, likely attributable to the overall increase of CD4^+^ T cells. Interestingly, of the infiltrating CD4^+^ T cells in the CNS, the frequency and number of CD4^+^RORγt^+^ cells were elevated **(Figure 1E**). Since T_H_17 cells can give rise to T_H_1-like cells that produce IFNγ, and acquisition of IFNγ (IL-17A^+^IFNγ^+^ cells) has been linked to pathogenicity in several models of chronic disease, including EAE/MS^12–14^, we assessed the frequency and number of IL-17A^+^, IL-17A^+^IFNγ^+^, and IFNγ^+^ T cells in the CNS (**Figure 1F**). While we did not observe a statistically significant increase in IL-17A^+^ T cells in the βKO relative to WT mice, there was a significant increase in the frequency and number of IL-17A^+^IFNγ^+^ population in βKO mice compared to WT controls. These results indicate that loss of REV-ERBβ leads to increased pro-inflammatory responses *in vivo* and that REV-ERBβ plays an important, non-redundant role to REV-ERBα in the development of EAE.

**Figure 1.**
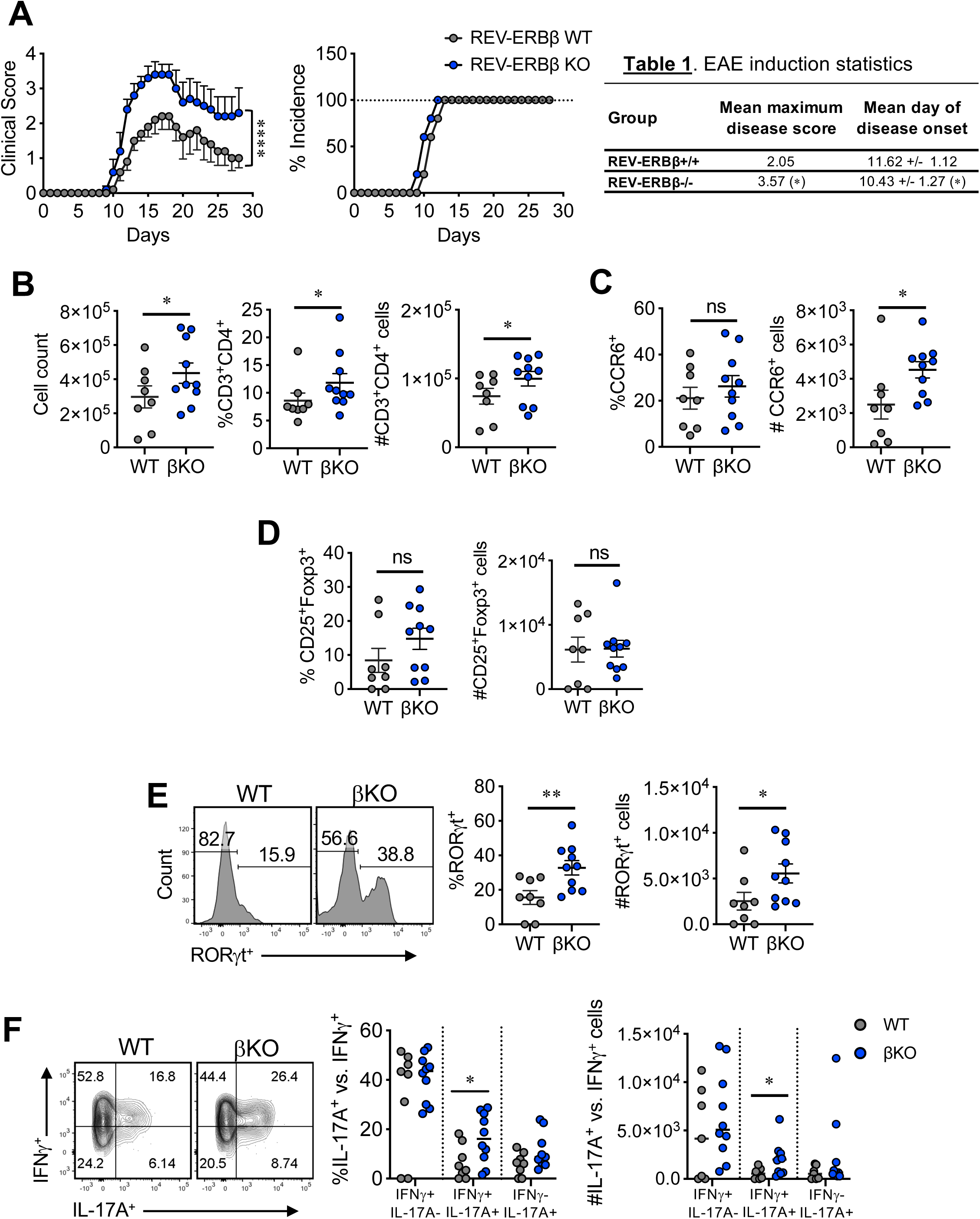
Loss of REV-ERBβ enhances the severity of EAE. **(A)** Clinical EAE scores (*left*) and disease incidence (*right)* of REV-ERBβ wild-type (WT) or knock-out (βKO) mice subjected to MOG-induced EAE. Table 1 (*right*) depicts mean maximum disease score and mean day of onset. **(B)** Graphs summarizing total cell count in CNS of WT vs. βKO mice (*left),* percent and number, respectively of CD3^+^CD4^+^ T cells at peak of disease (Day 15). **(C)** Graphs depict percent and number of CD4^+^CCR6^+^ cells in the CNS at peak of disease (Day 15). **(D)** Graphs depict percent and number of CD25^+^Foxp3^+^ cells in the CNS at peak of disease (Day 15). **(E)** FACS plots and graphs summarizing RORγt expression in the CNS of WT vs βKO mice at peak of disease. **(F)** FACS analysis and graph depicting the frequency and numbers of IL-17A^+^IFNγ^−^, IL-17A^+^IFNγ^+^, and IL-17A^−^IFNγ^+^ cells in the CNS of WT and βKO mice at peak of disease. For panels B-E, cells were gated on live, CD45^+^CD3^+^CD4^+^ cells. Each symbol represents an individual mouse. (n=8/WT, n=10/βKO). Data represents mean±s.e.m. Two-way ANOVA (clinical score) and Student’s *t*-tests were performed for statistical analysis. *\*p<0.05, **p<0.01, ****p<0.0001.* ns = not significant.

### REV-ERBβ-deficiency exacerbates the development of colitis

To determine the T-cell specific effects of REV-ERBβ *in vivo*, we sorted naïve CD4^+^ T cells from WT and βKO mice and adoptively transferred them into *Rag1^−/−^* recipients to track the development of colitis, which is another T_H_17-driven inflammatory disorder^13,15^, using PBS/sham injections as a control. Transfer of βKO cells resulted in more severe weight loss and increased colon weight (**Figure 2A, B**). Increased colon weight is a function of increased immune cell infiltration in the colon leading to exacerbated intestinal inflammation and disease. Measurement of colon length is also often used as a measure of disease, however, it is a delicate process and overzealous measuring techniques can skew results. To alleviate this bias and assess inflammation and edema in the colon, we assessed colon weight/length ratios. A higher ratio indicates more inflammation whereas a lower ratio suggests a healthier state. βKO mice presented with increased ratios relative to WT, indicative of more intestinal inflammation (**Figure 2B**). To further probe whether mice receiving βKO T cells presented with increased inflammation, we measured pro-inflammatory cytokine gene expression from proximal colon tissue. Expression of *Il17a*, *Il17f*, *Il6*, *Tnf*, and *Ifng*, was significantly increased in mice receiving βKO T cells compared to WT recipients when assessed by quantitative real-time polymerase chain reaction (qRT-PCR) (**Figure 2C**). Expression of *Il22* was also elevated in mice receiving βKO T cells but did not reach statistical significance (**Figure S2A)**. Finally, we observed increased expression of *Cxcl10* (**Figure S2C**), a gene that is activated downstream of IL-17 and IL-22 signaling, which is consistent with the overall increased T_H_17 proinflammatory profile observed *in vivo*.

**Figure 2.**
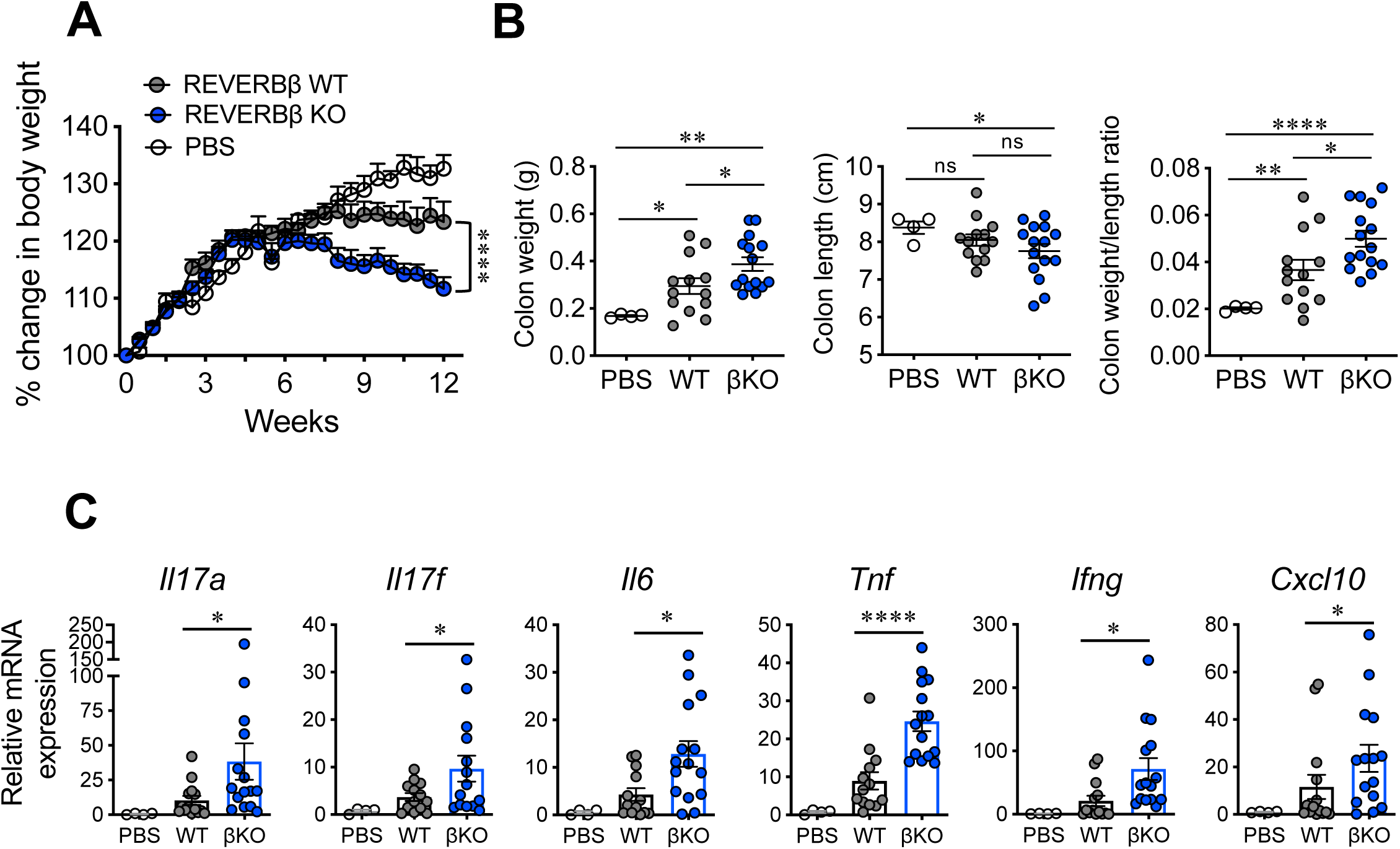
Loss of REV-ERBβ exacerbates colitis in mice. **(A)** Graph depicts percent change in body weight of *Rag1^−/−^*mice receiving REV-ERBβ wild-type (WT), knock-out (βKO) T cells, or control (no cells, PBS) for the long-term (12 week) experiment. **(B)** Graphs depict colon weights (*left*), colon lengths (middle), and colon weight/length ratio (*right*) of *Rag1^−/−^* mice receiving REV-ERBβ wild-type (WT), knock-out (βKO) T cells, or control (no cells, PBS) at the end of the 12-week experiment. **(C)** qRT-PCR analysis of pro-inflammatory cytokines and chemokines expressed in the (whole tissue) proximal colon from *Rag1^−/−^*mice receiving PBS, WT, or βKO CD4^+^ T cells. *18s* was used as the internal control. Each symbol represents an individual mouse. (n=13 for WT, n=15 for KO, and n=4 for PBS). Data represents mean±s.e.m. Two-way ANOVA (body weight) and Student’s *t*-tests were performed for statistical analysis. *\*p<0.05, **p<0.01, ****p<0.0001.* ns = not significant.

To specifically assess WT vs βKO T cell function, we performed similar experiments with the exception we sacrificed the *Rag1^−/−^* recipients at the peak of disease, 6-weeks post-transfer when T cells are starting to enter the colon. While we did not observe any changes in body weight between groups, we did observe increased colon weight in mice receiving βKO T cells relative to WT controls (**Figure 3A, S3A**). Colon weight/length ratios also indicated mice receiving βKO T cells presented with increased intestinal inflammation (**Figure 3A**). We also observed increased infiltration of total immune cells, frequency, and number of CD3^+^CD4^+^ T cells in the mesenteric lymph nodes (mLNs) and colons of *Rag1^−/−^* mice receiving βKO T cells vs. mice receiving WT cells (**Figure 3B, S3C**). No difference was observed in the spleen (**Figure S3B**). We also observed no differences in CD25^+^ Foxp3^+^ T regulatory cell frequencies and numbers in the colons between groups (**Figure S3D**). Further, while mice receiving βKO T cells did not have an increased frequency of CD4^+^RORγt^+^ or RORγt^+^IL-17A^+^ T_H_17 cells, there were increased numbers of CD4^+^RORγt^+^ and RORγt^+^IL-17A^+^ T_H_17 cells in the colons of recipient mice (**Figure 3C, D**). Finally, increased expression of pro-inflammatory cytokines and chemokines, including *Cxcl10*, *Ccr9*, and *Il22* was confirmed by NanoString experiments performed on FACS sorted T cells isolated from the colons of mice receiving βKO vs WT T cells (**Figure 3E**). This data is consistent with the increased gene expression observed in the colon tissues of βKO mice in the 12-week experiments suggesting increased IL-17 and IL-22 signaling (**Figure 2**). The increased expression of *Ccr9* could account for the increased infiltration of T cells to the colon as this chemokine receptor has been shown to facilitate T cell homing to the intestine under specific inflammatory conditions^16^. Interestingly, expression of genes involved in oxidative stress and apoptosis (both pro-and anti-apoptosis) were decreased in βKO T cells relative to WT controls, including *Sod1*, *Bcl2, Casp3, Casp8, Xiap, etc.* These data suggest REV-ERBβ may regulate apoptosis and cell survival in T cells in the colon. Further, expression of *Il18rap* was downregulated. This receptor has been shown to act as a negative regulator of T_H_17 cell differentiation in the intestine and its dysfunction or genetic variations are linked to intestinal diseases like Crohn’s disease, IBD, and celiac disease^17–19^. These results indicate that loss of REV-ERBβ promotes intestinal inflammation in our model of T-cell mediated colitis, leading to increased numbers of pro-inflammatory T_H_17 cells, reinforcing the role of REV-ERBβ as a negative regulator of T_H_17 responses.

**Figure 3.**
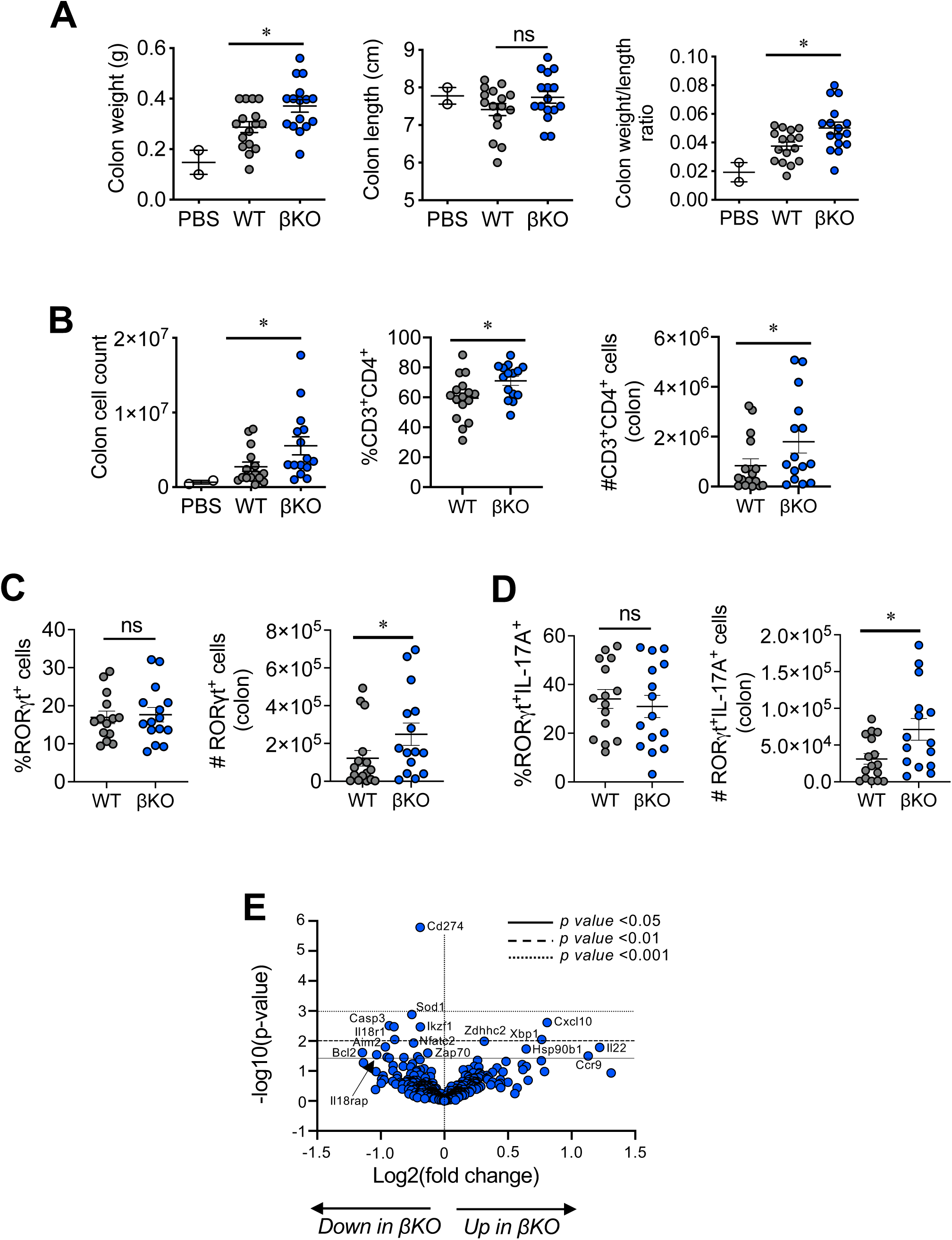
REV-ERBβ-deficient T cells promote colitis. **(A)** Graphs depict colon weights (*left*), colon lengths (*middle),* and colon weight/length ratios *(right*) of *Rag1^−/−^* mice receiving REV-ERBβ wild-type (WT), knock-out (βKO) T cells, or control (no cells, PBS) for the short-term (5-6 week) experiment. **(B)** Graphs summarizing total lymphocyte cell count (*left*), percent (*middle*), and number (*right*) of CD3^+^CD4^+^ cells in the colons of *Rag1^−/−^* recipient mice from the short-term experiments. Graphs summarizing frequencies, measured by FACS, and cell numbers of **(C)** RORγt^+^ and **(D)** RORγt^+^IL-17A^+^ cells in the colons of *Rag1^−/−^*recipient mice from the short-term experiments. Each symbol represents an individual mouse. (n=14 for WT, n=15 for βKO, and n=2 for PBS). **(E)** Nanostring data demonstrating differential gene expression changes between WT and βKO T cells isolated from the colons of *Rag1^−/−^*recipient mice 5-6 weeks post-transfer. Data is combined data from 4 donors each (WT vs. βKO). Data represents mean±s.e.m. Two-way ANOVA (body weight) and Student’s *t*-tests were performed for statistical analysis. *\*p<0.05, **p<0.01, ****p<0.0001.* ns = not significant.

### REV-ERBβ inhibits T_H_17 differentiation

To model REV-ERBβ-dependent T cell functions *in vitro*, we tested the requirement for REV-ERBβ to limit pro-inflammatory cytokine expression in CD4^+^ T helper cells. Using congenic co-cultures, we found that loss of REV-ERBβ decreased the ratio of βKO to WT T cells (CD45.1, WT; CD45.2, REV-ERBβ KO) when cultured under non-pathogenic (TGFβ + IL-6) and pathogenic conditions (pathogenic 1:TGFβ + IL-6 + IL-23; Pathogenic 2: IL-1β + IL-6 + IL-23) (**Figure 4A**). Loss of REV-ERBβ had no effect on the development of T_H_1 or inducible T regulatory (iTreg) cell development (**Figure S4**). Given we observed increased numbers of βKO cells in tissues in the T-cell transfer colitis model and EAE, these data suggest there may be a cell-extrinsic effect regulating their cell proliferation and/or survival. Despite the decreased frequency of βKO cells in the culture, REV-ERBβ-deficiency led to increased IL-17A production in T_H_17 cells when cultured under non-pathogenic or pathogenic 1 conditions but not pathogenic 2 conditions, suggesting a potential requirement for TGFβ in REV-ERBβ’s regulatory effects on T_H_17 cells (**Figure 4B**). To address the decrease in REV-ERBβ KO T cells in the co-culture, we stained the cells with carboxyfluorescein succinimidyl ester (CFSE) to assess proliferation and a viability dye to assess cell viability. Interestingly, while we did not observe any overt defects in proliferation, we did observe an increase in non-viable cells in culture with the βKO versus WT T cells, suggesting loss of REV-ERBβ lead to increased cell death in culture (**Figure 4C, D**). These data are consistent with the NanoString data in which REV-ERBβ KO T cells exhibited decreased expression of *Sod1*, *Bcl2, and Xiap*. These effects appear to be specific to T_H_17 cells as loss of REV-ERBβ did not affect the differentiation of T_H_1 or iTreg cells **(Figure S4**). These data suggest REV-ERBβ is required for downregulating effector responses in T_H_17 cells while maintaining cell viability.

**Figure 4.**
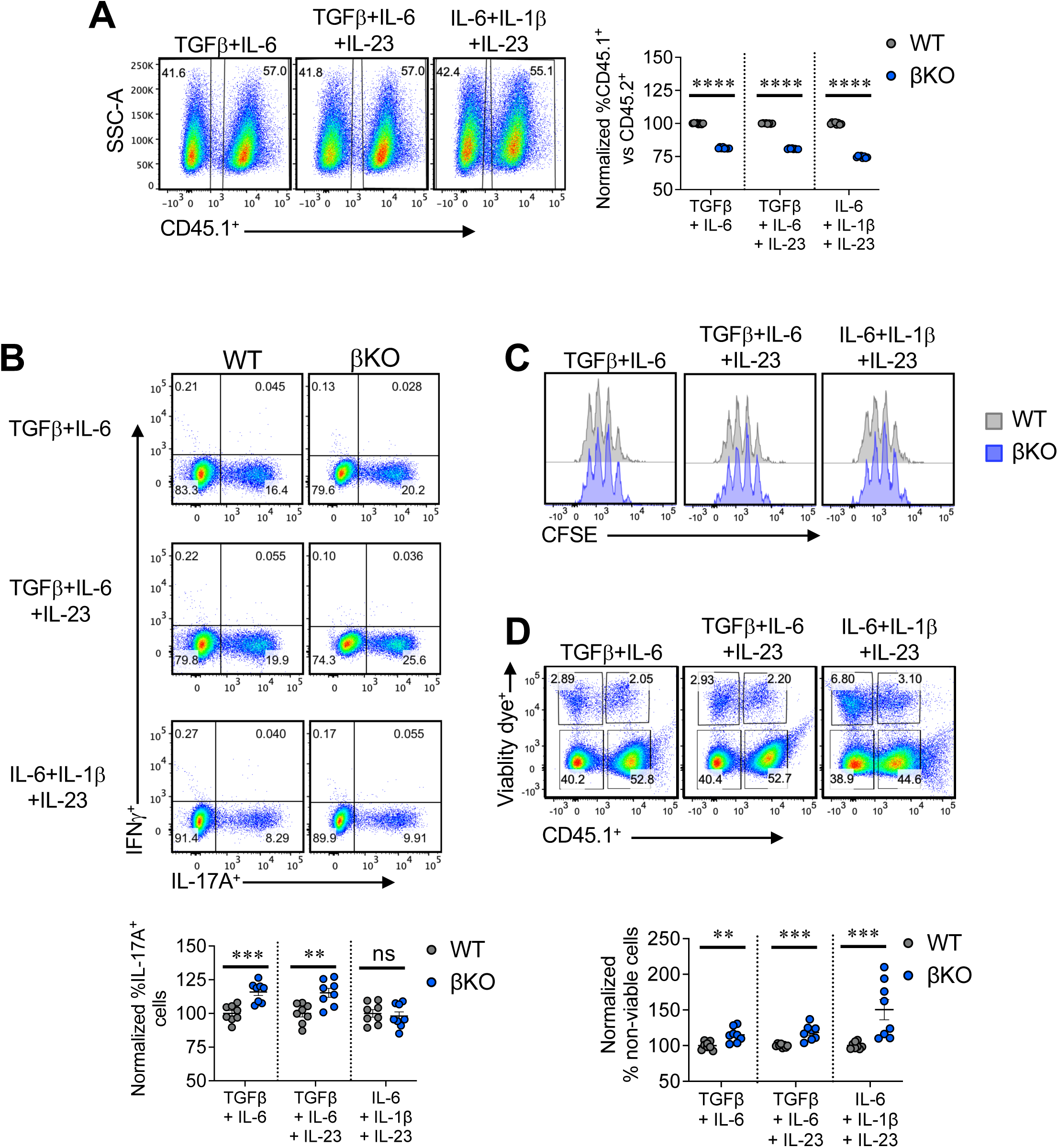
Cell intrinsic effects related to loss of REV-ERBβ in T cells. **(A)** Naïve CD4^+^ T cells from CD45.1 (WT) and REV-ERBβ T cell deficient (βKO) mice were differentiated under three different T_H_17 polarizing conditions (non-pathogenic; TGFβ+IL-6; pathogenic 1: TGFβ+IL-6+IL-23; pathogenic 2: IL-6+IL-1β+IL-23). Representative FACS plot from one experiment (*left panels*) demonstrating differences in percent CD45.1 cells. Right graph summarizing data demonstrating the frequency of WT vs. βKO T cells present in the cultures on day 4 post activation. **(B)** Representative FACS plots indicating the frequencies of IL-17A^+^ cells in the three different T_H_17 cultures (*top panels*). Bottom graph represents summary data from the different T_H_17 cultures. (**C**) CFSE data of WT vs βKO cells polarizing under the three different T_H_17 polarizing conditions. **(D)** Representative FACS plot from one experiment (*top panels*) demonstrating differences in percent CD45.1 cells and viability in each cell population. Bottom graph summarizing data of non-viable cells from three different T_H_17 polarizing conditions. Data are presented as mean values ±s.e.m. and representative of two-three combined independent experiments. Student’s *t*-tests were performed for statistical analysis. *\*\*p<0.01, ***p<0.001, ****p<0.0001.* ns = not significant.

### REV-ERBβ controls T_H_17 cell responses through a cell intrinsic manner

To address whether REV-ERBβ affects T_H_17 cell function through cell-intrinsic or cell-extrinsic mechanisms *in vivo*, we performed co-transfer experiments using the T-cell transfer model of colitis. We transferred equal numbers of congenic CD45.1^+^ (WT) or CD45.2^+^ REV-ERBβ KO (βKO) naïve CD4^+^ T cells (i.p.) into *Rag1^−/−^* recipients (**Figure 5A**). After 6 weeks of expansion, there were observable differences in the ratios between WT and βKO T cells in the mesenteric lymph nodes (mLN) and colon lamina propria of mice receiving WT vs KO T cells. These results corroborate our *in vitro* data. Effects in the spleen did not reach statistical significance (**Figure 5A**). Upregulation of the chemokine receptor CCR9 and the integrin α4β7 in mesenteric lymph nodes mediates lymphocyte homing to the intestinal lamina propria^20,21^. Evaluation of CCR9 and α4β7 expression indicated that while there was no difference in α4β7 expression in WT vs. βKO T cells, expression of CCR9 was increased in βKO T cells in the mLNs and colon (**Figure 5B**). Despite this, REV-ERBβ ΚΟ T cells did not have an advantage homing to the mLNs or colons in the donor mice. We also observed an increased frequency of RORγt^+^ cells and RORγt^+^IL-22^+^ cells in the βKO population relative to WT T cells (**Figure 5C,D**). IL-22 is a pro-inflammatory cytokine that influences intestinal barrier homeostasis^22^. This data corelates with our previous colitis experiments and NanoString analysis. Finally, there was an increased frequency of IL-17A^+^IFNγ^+^ cells in the βKO populations. These results demonstrate that REV-ERBβ is required to control T cell accumulation at inflammatory sites as well as temper downstream T_H_17 pro-inflammatory responses.

**Figure 5.**
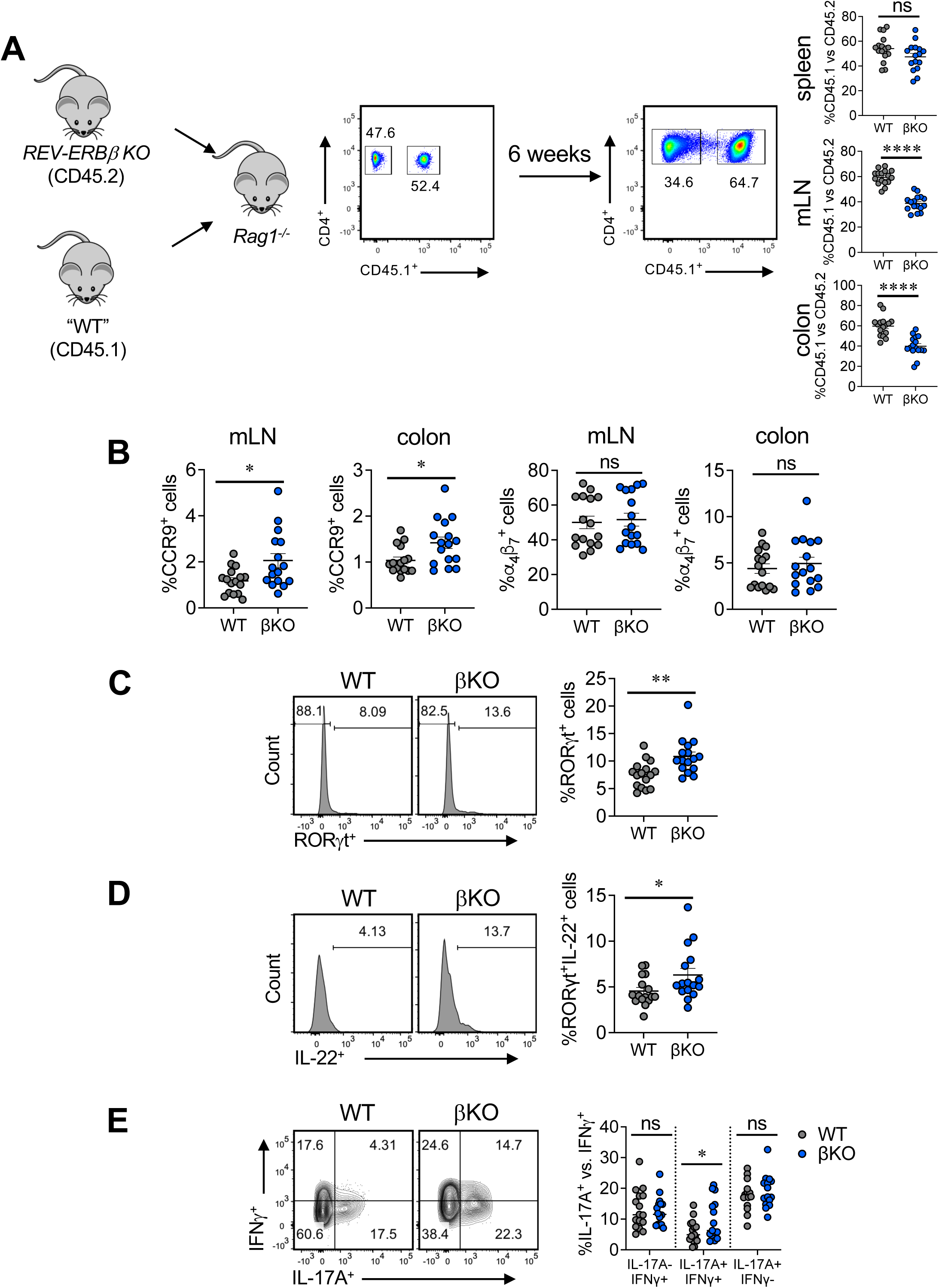
Loss of REV-ERBβ in T cells has cell intrinsic effects *in vivo* in colitis. **(A)** Schematic of experimental set up. Naïve CD4^+^ T cells were sorted from CD45.1 (WT) and REV-ERBβ T cell deficient (βKO) mice and transferred into *Rag1^−/−^*recipient mice. Representative FACS plot from one experiment (*left panel*). Right panels; representative FACS plots and graphs demonstrating the frequency of WT vs. βKO T cells present in the spleens, mLNs, and colons of *Rag1^−/−^* recipient mice 6 weeks post-transfer. (**B**) Graphs summarizing the frequency of CCR9^+^ and α4β7^+^ cells in mLN and colon of *Rag1^−/−^* recipient mice. FACS analysis and graphs depicting the frequency of (**C**) RORγt expression, **(D)** IL-22 expression, and (**E**) IL-17A^+^IFNγ^−^, IL-17A^+^IFNγ^+^, and IL-17A^−^IFNγ^+^ WT and βKO cells in the colons of *Rag1^−/−^*recipient mice at 6 weeks post-transfer. Cells were gated on live, CD3^+^CD44^+^CD4^+^ and CC45^+^ or CD45^−^ cells. Each symbol represents an individual mouse [n=16 (WT), n=16 βKO) for panels **Α-Ε**]. Data are presented as mean values ±s.e.m. Data are representative of two combined independent experiments. Student’s two-tailed *t*-tests were performed for statistical analysis. *\*p<0.05, **p<0.01, ****p<0.0001.* ns = not significant.

### REV-ERBβ regulates different genes than REV-ERBα in T_H_17 cells

We previously reported differential expression of REV-ERBα vs REV-ERBβ in T_H_17 cells and we wanted to verify this while comparing to RORγt’s expression^4^. Naïve CD4^+^ T cells were differentiated under T_H_17 polarizing conditions to assess the mRNA expression of REV-ERBβ (*Nr1d2*), REV-ERBα (*Nr1d1*), and RORγt (*Rorc*). qRT-PCR analysis demonstrated that REV-ERBα was upregulated similar to RORγt, however, REV-ERBβ was downregulated upon T_H_17 cell differentiation, similar to what we previously described^4^ (**Figure 6A**). This divergent gene expression pattern in T_H_17 cells between the REV-ERBs is in contrast to what is observed in other tissues, where the REV-ERBs typically have overlapping tissue expression patterns (i.e., liver, adipose tissue, brain, etc.)^7,8,23^. To gain a better understanding of the transcriptional program controlled by each receptor, we performed RNA-sequencing on T_H_17 cells collected at peak mRNA expression of each REV-ERB: time 0/naive for REV-ERBβ and 24h post-activation for REV-ERBα. We found that at both time points, there was very little overlap in transcriptional signatures between REV-ERBβ KO and REV-ERBα KO (αKO) T cells (**Figure 6B,C**). The genes that did overlap fell under “Metabolic pathways” via KEGG pathway analysis (i.e., *Rarg, Gcat, Abcb6, Slc43a1*, etc.). KEGG pathway analysis of cells at time 0 indicated REV-ERBβ regulated a large number of pathways relative to REV-ERBα, regulating genes involved in the cell cycle (i.e., *Ccnd2, Ccnd3, Ccnb1*, etc.), whereas REV-ERBα regulated genes involved in the circadian rhythm, which is consistent with its role as a core member of the circadian clock^23^. Additionally, REV-ERBβ also appeared to regulate genes involved in FoxO, JAK-STAT, and PI3K-Akt signaling pathways (i.e., *Il7r, Pik3r2, Ifngr1,* etc.) at time 0 (**Figure 6B**). Alternatively, in T_H_17 cells at 24h, REV-ERBβ regulated genes involved in Polycomb repressive complexes, Notch signaling pathways, regulation of the actin cytoskeleton, and cellular senescence, whereas REV-ERBα, as previously demonstrated, regulated genes involved in CD4^+^ T cell differentiation, including T_H_1, T_H_2, and importantly, T_H_17 cell differentiation (**Figure 6C**). Further, at 24h, REV-ERBα and REV-ERBβ again had very little overlap in transcriptional signatures, only appearing to collectively regulate genes for key transcription factors, cofactors, and chromatin remodeling factors (i.e., *Nfatc2, Nr4a3, Ncor2, Kmt2a*, etc.) (**Figure 6C**). At 24h, REV-ERBβ again regulated many cellular processes involved in negative transcriptional regulation as well as various signaling pathways that have been shown to be important regulators of T_H_17 cell development, including AMPK, FoxO, Notch signaling, and oxidative phosphorylation^24–26^. GO ontology analysis of the REV-ERBβ transcriptome revealed pathways involved in cell division at time 0 and chromatin and nucleosome organization at 24h post activation (**Figure 6D**). These data suggest that REV-ERBβ largely functions in an independent manner from REV-ERBα in T_H_17 cells, regulating pathways important for T_H_17 cell development, cell cycle, and chromatin remodeling.

**Figure 6.**
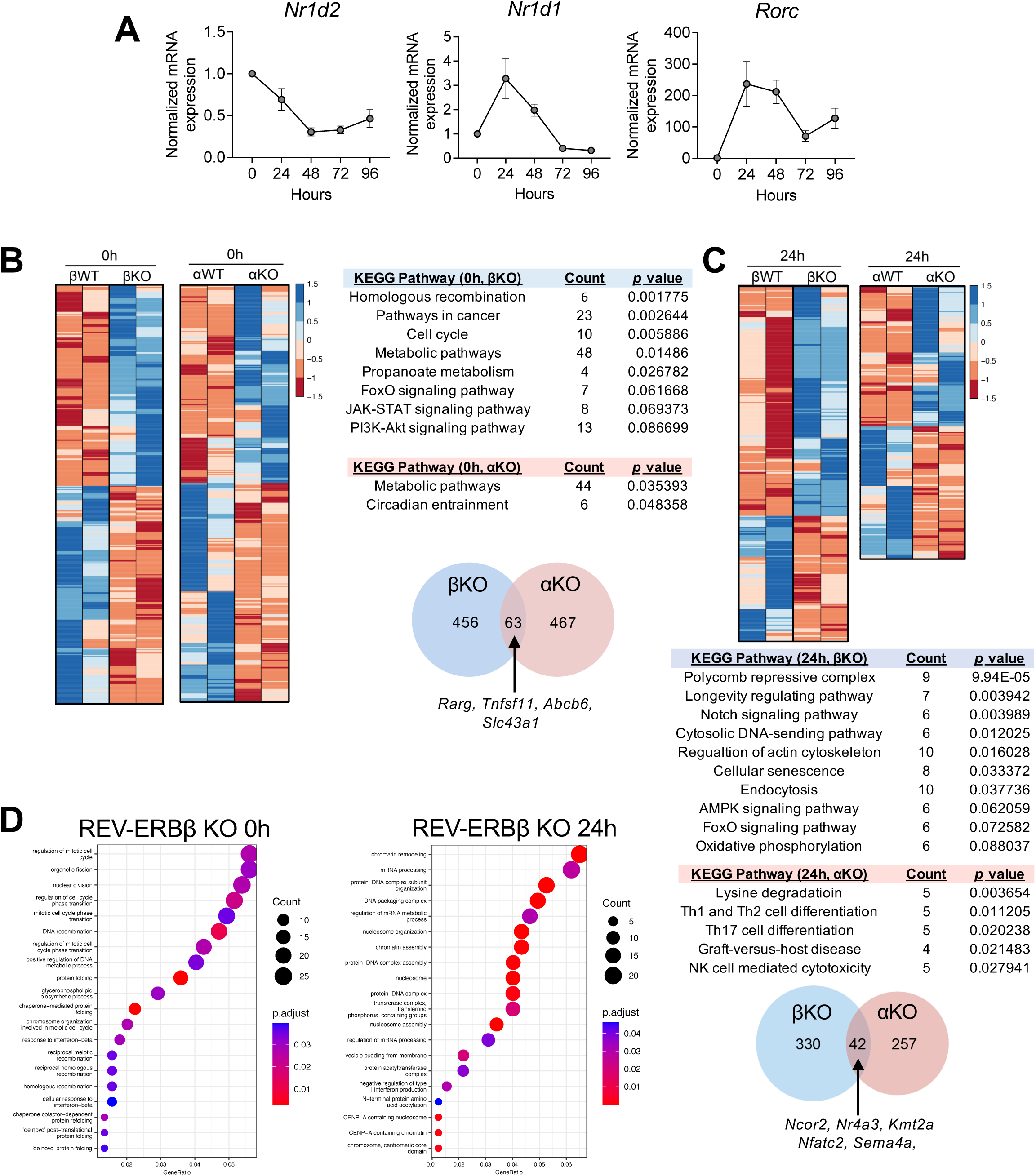
REV-ERBβ and REV-ERBα do not have overlapping gene programs. **(A)** qRT-PCR analysis of REV-ERBβ (*Nr1d2*), REV-ERBα (*Nr1d1*), and RORγt (*Rorc*) expression during T_H_17 cell development. β-actin was used as the internal control. **(B)** Heat map of differentially expressed genes in naïve CD4^+^ T cells from WT/REV-ERBβ ΚΟ vs. WT/REV-ERBα ΚΟ mice. KEGG pathway analysis of gene differentially expressed between the two receptors. Venn diagram (*bottom*) demonstrating overlap in transcriptional programs between βKO vs αKO cells. **(C)** Similar to panel B, with the exception that the comparison is between T_H_17 cells at 24h post activation. A small number of direct target genes are shared between the two receptors (FDR<0.05). **(D)** GO analysis of differentially expressed genes from βKO cells at time 0 and 24h post activation (T_H_17 cells). **(E)** REV-ERBβ LBD interacts with the NCoR corepressor. Co-transfection assay in HEK293 cells, which were transfected with the GAL4 UAS luciferase vector, GAL4-REV-ERBβ LBD, and an increasing concentration of NCoR VP-16 (or VP16 empty vector; EV). (*n=8*). Data are representative of 3 separate, independent experiments. Two-way ANOVA and Student’s *t*-tests were performed for statistical analysis. *\*\*\*\*p<0.0001.* **(F)** Venn diagram demonstrating overlap between NCoR ChIP and βKO differentially expressed genes in naïve CD4^+^ T cells. (FDR<0.05).

### REV-ERBβ does not appear to use NCoR1 for its transcriptional function

Transcription factor-DNA binding is one of the first steps towards modifications in gene transcription. Transcriptional cofactors need to be recruited, functioning as intermediaries between the transcription factors and the transcriptional machinery. A significant amount of work has been performed examining REV-ERBα and the cofactors it interacts with, identifying NCoR1, as the main corepressor instilling REV-ERBα with its repressive function^27^. Because of REV-ERBβ’s homology to REV-ERB*α*, and little interrogation of REV-ERBβ’s biological function, NCoR1 is also thought to be the prime corepressor responsible for both REV-ERBs’ repressive function^23,28^. To determine if REV-ERBβ interacts with NCoR1, we performed mammalian two-hybrid (M2H) protein-protein interaction assays in HEK293 cells co-transfected with three constructs: 1) REV-ERBβ ligand binding domain (LBD) which is fused to the yeast GAL4 DNA binding domain, 2) NCoR1 fused to the VP-16 activation domain, and 3) a UAS-luciferase reporter. When ligand binds, in this case, heme, the endogenous ligand for the REV-ERBs which is present in media^29,30^, a conformational change occurs in the REV-ERBβ LBD resulting in recruitment of NCoR1-VP-16 and increased luciferase signal^31^. Increasing concentrations of NCoR1-VP-16 drove increased luciferase signal, indicating a dose-dependent interaction with GAL4 REV-ERBβ LBD suggesting REV-ERBβ can interact with NCoR1, which is consistent with previously published data^31^ (**Figure 7A**).

**Figure 7.**
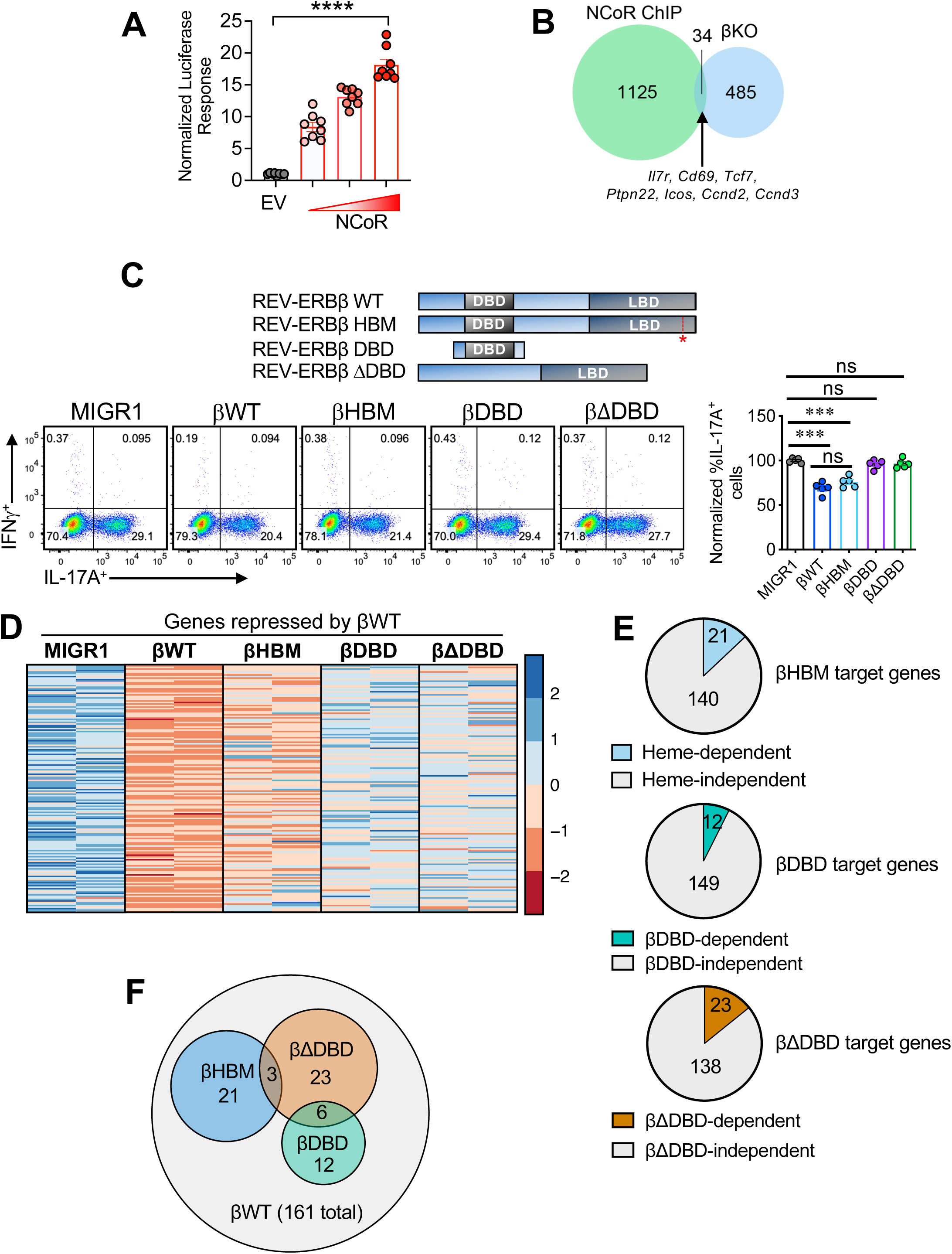
REV-ERBβ does not rely on NCOR1 recruitment for gene regulation. **(A)** REV-ERBβ LBD interacts with the NCoR corepressor. Co-transfection assay in HEK293 cells, which were transfected with the GAL4 UAS luciferase vector, GAL4-REV-ERBβ LBD, and an increasing concentration of NCoR VP-16 (or VP16 empty vector; EV). (*n=8*). Data are representative of 3 separate, independent experiments. Two-way ANOVA and Student’s *t*-tests were performed for statistical analysis. *\*\*\*\*p<0.0001.* **(B)** Venn diagram demonstrating overlap between NCoR ChIP and βKO differentially expressed genes in naïve CD4^+^ T cells. (FDR<0.05). **(C)** Schematic demonstrating REV-ERBβ constructs. Red (*) marks REV-ERBβ heme binding mutation (βHBM) - H565F (*top*). Representative FACS plots indicating the effects of the REV-ERBβ retroviral constructs on the frequencies of IL-17A in the different T_H_17 cultures (*bottom panels*). Graph represents summary data from the different T_H_17 cultures. **(D)** Heat map showing genes differentially repressed in the various REV-ERBβ T_H_17 conditions relative to empty vector (MIGR1). Genes are colored according to relative expression with normalization per gene across each row. (FDR<0.05). **(E)** Fraction of genes that are dependent on the different REV-ERBβ domains or constructs in T_H_17 cells relative to REV-ERBβ WT. **(F)** Venn diagram demonstrating the amount of overlap in transcriptional programs between constructs relative to REV-ERBβ WT. A small number of direct target genes are shared between the constructs, with the exception of REV-ERBβ WT. Two-way ANOVA and Student’s *t*-tests were performed for statistical analysis. *\*\*\*p<0.001, ****p<0.0001.* ns, not significant (*p>0.05*).

To determine if a REV-ERBβ-NCoR1 interaction regulates gene transcription in CD4^+^ T cells, we utilized a publicly available ChIP-seq data set in which NCoR1 binding sites in naïve CD4^+^ T cells were probed^32^. Notably, *Ncor1* is at its peak mRNA expression in naïve CD4^+^ T cells, much like REV-ERBβ (*Nr1d2*)^32^. Overlay of NCoR1 ChIP-seq peaks with differentially expressed genes from naïve CD4^+^ T cells (time 0) from the WT vs REV-ERBβ KO RNA-seq analysis revealed very little overlap between the two datasets with only 34 shared target genes (12 down-, 22-upregulated by REV-ERBβ) (**Figure 7B**). Interestingly, of the overlapping genes between NCoR1 and REV-ERBβ, many of them were T cell activation-dependent genes (i.e., *Cd69*, *Ctse*), and/or play significant roles in T cell development, proliferation, and function (i.e., *Ccnd2, Ccnd3, Icos, Il7r, Ptpn22*, etc.). These data indicate that the shared target genes of NCoR1 and REV-ERBβ are related more toward overall T cell activation and proliferation than T_H_17 lineage specification. This data is in line with NCoR1’s function in naïve CD4^+^ T cells, which suggests this co-repressor regulates general activation and proliferation genes rather than lineage-specific genes^32^.

Considering the outcome of the NCoR1 ChIP-seq/REV-ERBβ RNA-seq analysis, we next wanted to probe the molecular mechanisms of REV-ERBβ-dependent repression in T_H_17 cells. We and others previously showed that REV-ERBα represses T_H_17 cell differentiation by competing with RORγt for shared DNA binding sites within the IL-17A promoter^4,5^. The REV-ERBs are thought to repress transcriptional activity through either displacement of the RORs (“passive”) or recruitment of corepressor proteins, including NCoR1 (“active”), when bound to a RORE/RevRE DR2 DNA motif ^27,33^. To determine whether this occurs via passive or active repression with REV-ERBβ, we generated REV-ERBβ constructs encompassing the DBD alone (βDBD) or all but the DBD (βΔDBD) in the MIGR1 retroviral vector (**Figure 7C**). We also wanted to determine whether ligand regulation, which is thought to recruit co-factors, would affect REV-ERBβ’s transcriptional activity relative to other REV-ERBβ deletion constructs. As heme is the endogenous ligand for REV-ERBβ, and we recently identified the structural basis for heme-dependent NCoR1 recruitment, we asked whether heme is important for REV-ERBβ’s repressive activity^28^. However, it was previously shown that heme treatment represses T_H_17 cell differentiation by activating heme oxygenase (HO-1)^34,35^. To avoid REV-ERB-independent effects of adding heme to T_H_17 cell cultures, we introduced a point mutation in the ligand binding domain (LBD) in which the axial histidine critical for coordinating the central iron atom within the heme molecule is mutated to phenylalanine. This REV-ERBβ heme binding mutation (βHBM) was previously shown to reduce heme binding affinity >10,000-fold for REV-ERBβ (H565F)^29,36^. When we retrovirally overexpressed these constructs in T_H_17 cells and gated on transduced (GFP^+^) cells, we found the truncation and deletion constructs (ΒDBD and βΔDBD) were unable to repress *Il17a* relative to WT REV-ERBβ (βWT) (**Figure 7C**). The requirement for both the DBD and the LBD for transcriptional repression indicates REV-ERBβ exerts active repression by binding DNA (via its DBD) and recruiting corepressor (via its LBD). Interestingly, the βHBM did not relieve repression at IL-17A relative to the βWT construct (**Figure 7C**), indicating heme is not required for regulation of REV-ERBβ-mediated IL-17A repression.

We next performed RNA-sequencing of T_H_17 cells overexpressing these constructs to determine how they affected REV-ERBβ-mediated transcription. Similar to our FACS data, we found the βHBM had little effect on transcription, relieving repression of ∼10-15% of βWT target genes, confirming that heme is not very important for REV-ERBβ activity in T_H_17 cells (**Figure 7D,E**). Similarly, the βDBD and βΔDBD constructs did not regulate very many genes, again suggesting that REV-ERBβ exerts active repression by binding DNA and recruiting corepressor via its LBD. However, recruitment of co-repressor does not appear to be heme-dependent. Further, each mutant construct regulated transcription associated with βWT-dependent regulation but largely had their own transcriptional signatures as very little overlap was observed in the differentially expressed genes between the βHBM, βDBD, and βΔDBD (**Figure 7F**). Collectively, this data demonstrates that REV-ERBβ functions in a heme-independent manner in T_H_17 cells and use of its DBD and LBD appear to be required for its repressive function.

## DISCUSSION

In this study we demonstrate a non-redundant role for REV-ERBβ in the regulation of T_H_17 cell development and disease pathogenesis compared to REV-ERBα. We show that *in vitro*, loss of REV-ERBβ led to enhanced expression of IL-17A, but at the expense of cell viability. This did not extend to other T helper lineages evaluated, including T_H_1 and Foxp3^+^ T regulatory cells. Using a mouse model of EAE we show that loss of REV-ERBβ enhanced disease development and the expression of IL-17A and RORγt *in vivo*. In a mouse model of colitis, *Rag1^−/^*^−^ mice receiving REV-ERBβ KO T cells developed exacerbated signs of disease, due to increased numbers of T cells homing to the intestines. This effect was cell intrinsic as co-transfer studies indicated that βKO T cells exhibited increased expression of IL-22, IL-17A, and RORγt. Further, these effects appear to be distinct to REV-ERBβ as RNA-sequencing of REV-ERBα vs. REV-ERBβ KO CD4^+^ T cells indicated very little overlap in gene expression signatures between the two nuclear receptors. Finally, while REV-ERBβ’s repressive function appears to largely be independent of the co-repressor NCoR1 and heme, full-repression does still appear to be dependent on its ability to bind DNA and recruit co-repressor via its LBD.

Many in the nuclear receptor field believe that REV-ERBα and REV-ERBβ are functionally redundant, with REV-ERBα acting as the dominant factor^7^. However, prior data from our lab demonstrated that REV-ERBβ overexpression in T_H_17 cells did not compensate for REV-ERBα^4^. Our studies using REV-ERBβ knock out mice and subsequent RNA-sequencing experiments support this as comparison of the transcriptomes between the two receptors at different time points yields very little overlap in gene expression profiles. Thus, at least in the case of T_H_17 cells, there appears to be little redundancy between the REV-ERBs. In fact, given the different tissue pattern expressions, it would appear that the two receptors work in concert to suppress gene expression at different stages of T_H_17 cell development to keep aberrant pro-inflammatory responses in check.

Molecular, biochemical, and cell-based evidence suggest that both REV-ERBα and REV-ERBβ require their endogenous ligand, heme, to interact with NCoR1 to repress transcription^28–30,36^. This REV-ERBα-NCoR1 interaction has been shown to occur at *Il17a* to repress its expression^5^. Importantly, the REV-ERBα-NCoR1 interaction has been shown to then recruit the histone deacetylase, HDAC3, to further repress transcription^37^. However, our data indicates that, at least in T_H_17 cells, REV-ERBβ mediates its repression independently of heme and NCoR1, which is in contrast to REV-ERBα and provides further evidence that REV-ERB function is divergent in this cell type. In fact, much of the work interrogating REV-ERB/co-factor interaction has been via assessment of REV-ERBα with further assumptions that REV-ERBβ functions similarly due to its “redundant” nature to REV-ERBα. However, one publication from 2008 interrogated REV-ERBβ’s interacting partners indicating this may not be the case. In this publication, REV-ERBβ recruited the histone acetyl-transferase, Tip60, via its N-terminus in a ligand-independent fashion, to the *ApocIII* promoter to promote its transcription, and recruited the histone deacetylase, HDAC1, via its LBD, to repress gene transcription^38^. This publication provided evidence that REV-ERBβ can modify gene expression in a positive and negative manner through different co-factor interaction than REV-ERBα. Given the small number of genes that were regulated by both NCoR1 and REV-ERBβ, the number of genes that were upregulated and downregulated by REV-ERBβ deficiency, and that the heme binding mutant failed to relieve REV-ERBβ-mediated repression, it is conceivable that other non-canonical cofactors, similar to Tip60 and HDAC1, are mediating REV-ERBβs transcriptional effects. Thus, more work needs to be performed to explore the cofactors and mechanism of action by which REV-ERBβ mediates transcriptional repression.

Although we previously showed that the REV-ERBs’ repressive activity in T_H_17 cells is enhanced by synthetic small molecules, the role of the natural REV-ERB ligand, heme, remained to be defined. Our findings incorporating REV-ERBβ truncation and deletion constructs showed that REV-ERBβ requires all intact domains for complete repression, suggesting it represses by active mechanisms involving DNA and coregulator binding. Given heme has previously been shown to be important for active repression^28^, we explored whether heme is important for REV-ERBβ activity in T_Η_17 cells using a point mutation that specifically disrupts heme binding. These experiments showed heme is not important for full REV-ERBβ-mediated repression of target genes in T_H_17 cells *in vitro*. Collectively, our results suggest a non-canonical signaling mechanism for REV-ERBβ in which heme is not critical for its repressive function.

To date, several small molecules targeting both REV-ERBα and REV-ERBβ have been developed as an alternative to RORγt modulators^4–6^. We, and others, have previously published these small molecules are potent inhibitors of T_H_17-mediated disease processes without the negative effects on the thymus observed with RORγ inhibitors. One ligand we recently developed is even selective for REV-ERBα over REV-ERBβ^6^. However, our current data suggests that targeting both factors would be ideal to better encompass inhibition of the T_H_17 program *in vivo*, both during the initial development of T_H_17 cells and later post activation, when REV-ERBα is more highly expressed. By this logic, it would stand to reason that ligands that target both REV-ERBs would have a greater repressive effect than a ligand that targeted only a single receptor subtype in T_H_17 cells.

In summary, we demonstrate that the nuclear receptor REV-ERBβ acts in a non-redundant manner to REV-ERBα during T_H_17 cell development *in vitro* and *in vivo*. While there is still much work to be done to fully elucidate the function of both REV-ERBs in T_H_17 cells, our work demonstrates the potential for individuality between receptors, something that has never been shown before. Thus, our data suggest that REV-ERBβ may not be as functionally redundant to REV-ERBα as originally hypothesized.

## RESOURCE AVAILABILITY

### Lead contact

Requests for further information and resources should be directed to and will be fulfilled by the lead contact, Laura A. Solt.

### Materials availability

All unique reagents generated in this study will be made available upon request.

### Data code and availability

- All RNA-sequencing data can be made available upon request to the lead contact.
- This paper analyzes existing, publicly available data, accessible at GSE138934.
- This paper does not report original code.
- Any additional information required to reanalyze the data reported in this paper is available from the lead contact upon request.

## ACKNOWLEDGEMENTS

We thank the UF Scripps Genomics and Bioinformatics Cores for library preparation, RNA sequencing, and data analysis. This work was supported by the US National Institutes of Health (R01AI116885, R21AI164722, and R01CA241816 to L.A.S. and F31DK127643 to S.A.M.). M.A. was supported through the American Association of Immunologists Careers in Immunology Fellowship Program and the Crohn’s and Colitis Foundation of America (#546172).

## AUTHOR CONTRIBUTIONS

L.A.S. conceived the project. L.A.S., M.A., S.A.M., Q.L., J.C., and S.C., designed/analyzed and/or performed the mouse *in vitro* assays and *in vivo* studies. L.A.S. wrote the manuscript with edits from all authors.

## DECLARATION OF GENERATIVE AI AND AI-ASSISTED TECHNOLOGIES

The authors declare that no Generative AI was used in creation of this manuscript.

## SUPPLEMENTAL INFORMATION

Document S1. Figures S1-S4

## METHODS

### Mice

The following mouse strains used were purchased from the Jackson Laboratory and/or were bred at UF Scripps Biomedical Research. C57BL/6J (B6); B6.129S7-*Rag1^tm1Mom^*/J (*Rag1^−/^*^−^); B6.SJL-*Ptprc^a^Pepc^b^*/BoyJ (CD45.1); B6.Cg-*Nr1d1^tm1Ven^*/LazJ (REV-ERBα^−/−^). *Nr1d2*-deficient mice have been previously described^39^. All experiments were conducted at controlled temperature (22–23°C), humidity ∼60%, and 12h:12h light:dark cycles. Mice had access to regular chow (Harlan 2920X) and water, *ad libitum.* For all *in vitro* experiments, both male and female mice (8-10 weeks old) were sacrificed between 8 and 10am. For all *in vivo* experiments, 7-10 week-old male and female mice were used, and were sacrificed between 7 and 11am. For EAE experiments, mice were immunized between 11am and 1pm. The specific age and sex of the mice for experiments is described under each model’s details below. All mice were maintained under specific pathogen free conditions. All studies conform to and were approved by the Institutional Animal Care and Use Committee (IACUC) at UF Scripps.

### Cell culture and maintenance

HEK293 cells and PlatE^40^ cells were cultured in DMEM supplemented with 10% FBS, 2mM L-glutamine, and 1% penicillin/streptomycin at 37°C, 5% CO_2_ under standard culture conditions. Lymphocytes were cultured in IMDM medium with 10% FBS, 100IU/mL penicillin, 100 μg/ml streptomycin, 50uM β-mercaptoethanol, and 2mM L-glutamine.

### Mouse *in vitro* CD4^+^ T cell differentiation

Naïve CD4^+^ T cells from spleen and lymph nodes of male and female 8-10-week-old mice were purified after removing the red blood cells using Lympholyte-M solution (Cedarlane Laboratories). Cells were enriched for naïve CD4^+^ T cells using the mouse naïve CD4^+^ T Cell Isolation Kit (STEMCELL Technologies, Canada) according to the manufacturer’s instruction. If sorting was performed (FACS Aria II; BD Bioscience), the CD4^+^CD25^−^CD62L^hi^CD44^lo^ fraction was collected. The conditions for the different T_H_ cell subsets were: For T_H_1 conditions: 5 μg/ml anti-IL-4, 20 ng/ml IL-12 (R&D Systems), and 10ng/ml IFNγ (R&D Systems); For T_H_17 conditions: (non-pathogenic) 5 μg/ml anti-IFNγ, 5 μg/ml anti-IL-4 (R&D Systems), and 1.5ng/ml TGFβ (R&D Systems) and 30ng/ml IL-6 (R&D Systems); (pathogenic 1) 5 μg/ml anti-IFNγ, 5 μg/ml anti-IL-4 (R&D Systems), and 1.5ng/ml TGFβ (R&D Systems), 30ng/ml IL-6 (R&D Systems), and 20ng/mL IL-23 (R&D Systems); (pathogenic 2) 5 μg/ml anti-IFNγ, 5 μg/ml anti-IL-4 (R&D Systems), and 30ng/ml IL-6 (R&D Systems), 10ng/ml IL-1β (R&D Systems) and 20ng/mL IL-23 (R&D Systems); For iT_reg_ conditions: 5 μg/ml anti-IFNγ, 5 μg/ml anti-IL-4, and 5ng/ml TGFβ (R&D Systems). 1 × 10^6^ cells/ml of naïve CD4^+^ T cells were activated with anti-CD3 (clone 2C11, 0.5 μg/ml) and anti-CD28 (clone 37.51; 1 μg/ml) by precoating plates with 50 μg/ml goat anti-hamster IgG. For congenic co-culture experiments, 5 × 10^5^ cells/ml of naïve CD45.1 CD4^+^ T cells and 5 × 10^5^ cells/ml of naïve CD45.2 CD4^+^ T cells were activated mixed together with anti-CD3 and anti-CD28, as described above. After 48 hr., cells were removed from the TCR signal and recultured at a concentration of 1 × 10^6^cells/ml. Four days after activation, all cells were restimulated with 50ng/mL phorbol-12-myristate-13-acetate (PMA) (Sigma) and 1 μg/ml ionomycin (Sigma) for 2 hours with the addition of GolgiStop (BD Bioscience) for an additional 2 hours before intracellular staining. Cells were cultured in IMDM medium (Invitrogen) with 10% FBS, 100IU/mL penicillin, 100 μg/ml streptomycin, 50uM β-mercaptoethanol, and 2mM L-glutamine. All cultures were performed in a volume of 200θμl in 96-well U-bottomed plates.

### Induction and clinical evaluation of EAE

EAE was induced in 10-week-old, male and female *Nr1d2*-deficient or wild-type mice using EAE induction kits (EK-2110; Hooke Laboratories, Lawrence, MA, USA) according to manufacturer’s instructions and as previously described^4,41^. Briefly, mice were injected subcutaneously over two sites in the flank with 200μg/mouse of mouse MOG_35-55_ peptide in an emulsion of Complete Freund’s Adjuvant (CFA), containing killed *M.tuberculosis*, strain H37Ra. Concentration of killed *M.tuberculosis* was adjusted by lot depending on EAE induction and ranged from ∼2-5mg/mL emulsion. Pertussis toxin was dissolved in PBS and injected i.p. at a range of 200-400ng/mouse 4 hours post immunization (day 0) and 24 hours later. The concentration of pertussis toxin used varied by lot and was determined by dose response studies. Clinical scoring started 7 days post immunization after which mice were scored daily according to the following criteria: 0, no clinical disease; 1, limp/flaccid tail; 2, limp tail and hind leg weakness; 3, limp tail and complete paralysis of hind limbs; 4, limp tail, complete hind limb and partial front limb paralysis; 5, quadriplegia or pre-moribund state. Gradations of 0.5 were used when mice exhibited signs that fell between two scores. All scoring was blinded to genotype and previous scores for each mouse. Disease was monitored for 28 days or as indicated in the figure legend. At the end of the experiment, mice were anaesthetized and transcardially perfused with PBS. The spleen, peripheral lymph nodes together with spinal cords were removed for single cell isolation for FACS analysis.

### T-cell Transfer Model of Colitis

Spleens were collected from 8-10-week-old, female *Nr1d2^+/+^* or *Nr1d2^−/−^*mice to sort naïve CD4^+^ T cells. A total of 5 × 10^5^ CD4^+^CD25^−^CD62L^hi^CD44^lo^ cells suspended in PBS were adoptively transferred i.p. (100μl/mouse) into 8-week old, female *Rag1^−/−^* recipient mice^42^. For congenic transfers, similar protocols were used with the exception that naïve female CD45.1 CD4^+^ T cells were sorted in lieu of *Nr1d2^+/+^* T cells. Post sort, cells were counted, adjusted for cell number, mixed at a 50/50 ratio, and analyzed by flow cytometry to determine equal frequency of the two experimental groups. Once verified, the mixture was injected into female *Rag1^−/−^* recipients. Mice were monitored bi-weekly for body weight change. Beddings were transferred between cages to minimize the impact of microflora on disease development. Mice were sacrificed to assess mRNA expression of inflammatory genes or due to ethical requirements if they reached 80% or less of their original body weight.

### Mouse intestine tissue end-point collection

At the termination of the 12-week experiments, the whole colon and ileum (distal 1/3 part of the small intestine) were removed from the mouse. Each segment of the intestine was opened longitudinally, and the fecal contents were gently removed. Colon length and weight were measured and then the colon was dissected in equal halves, designated as ‘proximal’ and ‘distal’ colon and snap frozen in dry ice for protein and RNA analysis.

### Lamina propria mononuclear cell Isolation from mouse intestine

At the termination of the 4-6-week experiments, to collect mononuclear cells from the lamina propria of intestine, mouse ileum or colon were dissected, opened longitudinally, rinsed three times in cold PBS, and then in DMEM phenol-free medium with 200 U/mL penicillin and 200 mg/mL streptomycin one time to remove the fecal content. Washed intestines were incubated in DMEM with 1 mM DTT at room temperature (RT) on a rocker for 30 min then in DMEM with 0.5M EDTA for 30 min at RT to remove the mucus layer. After washing twice in DMEM, intestines were then digested in DMEM with Liberase and DNase I at 37 C in a shaking water bath for 30 min. The cells were then dislodged from the lamina propria and submucosa by vigorous shaking for 30 seconds. The supernatant containing leucocytes was pooled together by centrifugation at 500g for 5 min. Cells were further cleaned up by 30%/70% Percoll (Sigma) gradient separation before further study.

### Flow cytometry

<u>Surface staining</u>: single cell suspensions prepared from spleen, lymph nodes, CNS, etc. were washed and stained with fluorescence-conjugated antibodies for 20 minutes, washed, then resuspended in FACS buffer (0.5% BSA, 2mM EDTA in PBS). <u>Intracellular cytokine staining</u>: cells were re-stimulated with 50ng/mL PMA and 1 μg/ml ionomycin for 2 hours with the addition of GolgiStop (BD Bioscience) for an additional 2 hours. Cells were then surface stained using procedures outlined above, fixed and permeabilized using the Foxp3 staining kit (eBioscience). Flow cytometric analysis was performed on a BD LSRII (BD Biosciences) instrument and analyzed using FlowJo software. All antibodies used in experiments are described in Key Resources.

### Retroviral Transduction

The MIGR1 vectors containing murine REV-ERBβ^4^ was used to generate mutant and truncation constructs. The DBD truncation construct was generated by PCR amplification of residues 73–195 of REV-ERBβ with addition of 5’ XhoI and 3’ HpaI cut sites. The amplified product was cloned into the MIGR1 vector by double digestion followed by T4 DNA ligation. The ΔDBD deletion construct was generated by a single PCR reaction to delete residues 102–167 of the MIGR1 REV-ERBβ construct. The H565F REV-ERBβ mutant was generated using site-directed mutagenesis. <u>Virus production</u>: Plat-E cells (Cell Biolabs, Inc.) were cultured in DMEM containing 10% fetal bovine serum, 2mM L-glutamine, and 1% penicillin/streptomycin at 37°C under standard culture conditions. Plat-E cells were seeded at 350,000 cells/ml in a 6 well plate the day before transfection. 3μg total retroviral plasmid DNA (1.5 μg MIGR1 plus 1.5 μg pCL-Eco) was transfected using Fugene6 reagent (Promega) according to manufacturer’s protocol. Viral supernatant was harvested 48 hours post transfection and used immediately for transduction. For <u>retroviral transduction</u>, naïve CD4^+^ T cells were stimulated as indicated with anti-CD3 and anti-CD28 and cultured under T_H_17 conditions. At 24 h post TCR priming, the culture medium was replaced with virus supplemented with 8 μg/ml polybrene. Plates were centrifuged at 1,800 rpm for 90 min at 37°C and then incubated at 37 °C for 3-4 h. After this time, the medium was replaced with the original media removed before addition of virus.

### Measurement of proliferation using Cell Trace Violet

Cell Trace Violet (CTV) was diluted to 10μM in room temperature PBS. Naïve T cells to be stained were resuspended in at a concentration of 20 × 10^6^ cells/ml in sterile room temperature PBS. Cells were gently vortexed while an equal volume of CTV dye was added into the cell suspension. The cell suspension was then vortexed for an additional 15 seconds and incubated in the 37°C water bath for 2.5 minutes CTV staining was immediately quenched with 4mls ice cold 1:1 PBS/FBS. Cells were pelleted at 2000rpm for 3 minutes and washed again with 4mls ice cold 1:1 PBS/FBS. T cells were pelleted and resuspended T cell media and plated as described above.

### FACS sorting of naïve CD4^+^ T cells

All FACS sorting was performed using a FACS Aria II (BD). Sorting of naïve CD4^+^ T cells: For *Rag1*^−/−^ transfer experiments, magnetically enriched CD4^+^CD25^−^ T cells (STEMCELL Technologies), obtained from spleens and lymph nodes, were FACS-sorted to obtain pure naive T cells (CD3^+^CD4^+^CD25^−^CD62L^hi^CD44^lo^). Similar methods were used for sorting naïve CD4^+^ T cells for RNA-sequencing.

### Quantitative Reverse-Transcription Polymerase Chain Reaction

Tissue samples (snap frozen and homogenized) were extracted into TRIzol before RNA purification using RNeasy columns (Zymo Research, CA), followed by cDNA synthesis using iScript (BioRad, CA) containing oligo (dT) and random hexamer primers. qPCR SYBR Green (Roche) was used for quantitative polymerase chain reaction using a HT7900 machine (Life Technologies, CA); passive reference dye ROX was used. Primer efficiencies were determined using complementary DNA and primer dilutions for each gene of interest. Primers used to determine gene expressions can be found in **Supplementary Table 1**. All gene expression data were normalized to the housekeeping gene *β-actin* or *18s*.

### Nanostring

Gene expression was quantified using the Mouse Autoimmunity panel (NanoString Technologies), containing 750 probe pairs with additional probes for positive and negative controls as well as housekeeping genes. Briefly, 20,000 T cells were lysed in 5μl of freshly made buffer RLT (Qiagen) containing β-ME. Biotin-conjugated capture probes and fluorescent-barcoded reporter probes were hybridized to cell lysates overnight at 65°C in a thermocycler. The following day, post-hybridization processing occurred on all lysates which were then run on an nCounter *SPRINT* (Nanostring Technologies) according to the manufacturer’s instructions. Raw data were normalized and analyzed using nSolver software (Nanostring Technologies).

### RNA-sequencing and data analysis

mRNA was extracted from naïve CD4^+^ T cells or T_H_17 cells 24h post activation (WT vs. REV-ERBβ KO cells or WT vs. REV-ERBα KO cells). Total RNA was extracted using Qiagen RNeasy kits, quantified using the Qubit 2.0 Fluorometer (Invitrogen, Carlsbad, CA), and run on the Agilent 2100 Bioanalyzer (Agilent Technologies, Santa Clara, CA) for quality assessment. DNase-treated total RNA (300ng) was depleted of ribosomal RNA (rRNA) using appropriate probes provided by Illumina (TruSeq Total RNA-seq kit) and further assessed on the bioanalyzer to confirm 18S and 28S rRNA peaks are depleted. rRNA-depleted RNA was processed using the TruSeq Stranded Total RNA sample prep kit (Illumina, San Diego, CA). Briefly, RNA samples were chemically fragmented in a buffer containing divalent cations and heated at 94°C for 8 minutes. The fragmented RNA was random hexamer primed and reverse transcribed to generate first strand cDNA. The second strand was synthesized after removing the RNA template and incorporating dUTP in place of dTTP. cDNA was then end repaired and adenylated at their 3’ ends. A corresponding ‘T’ nucleotide on the adaptors was utilized for ligating the adaptor sequences to the cDNA. The adaptor ligated DNA was purified using magnetic Ampure XP beads and PCR amplified using 12-13 cycles to generate the final libraries. The final libraries were size selected and purified using 1.0 x Ampure XP beads to remove any primer dimers. The final library size is/was typically 200-600bp with insert sizes ranging from 80-450bp. Final libraries were validated using bioanalyzer DNA chips and qPCR quantified using primers recognizing the Illumina adaptors. Libraries were pooled at equimolar ratios, quantified using qPCR (quantification of only the adaptor-ligated libraries) and loaded onto the NextSeq 500 flow cell at 1.8pM final concentration for pair end 75bp reads. 20-25 million mappable reads per sample were collected. Demultiplexed and quality filtered raw reads (fastq) generated from the NextSeq 500 were trimmed (adaptor sequences) using Flexbar 2.4 and aligned to the reference genome using TopHat version 2.0.9 ^43^. HT seq-count version 0.6.1 was used to generate gene counts and differential gene expression analysis was performed using DESeq2 ^44^. The normalized gene counts were used to plot the heatmaps. To determine enriched functional groups in the RNA-seq data, gene ontology (GO)^45,46^ enrichment analysis was performed using Cluster Profiler^47^ as was KEGG pathway analysis using DAVID^48,49^.

### ChIP-seq analysis

Target reads were mapped to the mm39 genome using Bowtie2^50^. Peaks were called over the control sample using MACS2^51^. ChIPseeker^52^ was used to annotate peaks to genes and GO enrichment analyses was completed^45,46^.

### Mammalian two-hybrid reporter assays

HEK293 cells were plated 24 hours prior to transfection in 96-well flat-bottom plates at a density of 15 × 10^3^ cells/well. Cells were co-transfected with 100 ng pG5-UAS vector and 25 ng pCMV-Gal4-REV-ERBβ LBD, along with pACT empty vector (Promega) expressing the VP16 transactivation domain only or increasing concentrations of pACT with VP16 fused to the mouse NCoR receptor interaction domain (RID, residues 1828–2471). Transfections were performed using Lipofectamine 3000 (Invitrogen) according to manufacturer’s protocol. Twenty-four hours post transfection, luciferase activity was measured using BriteLite (PerkinElmer Life and Analytical Sciences) and read using BioTek Synergy plate reader (PerkinElmer Life and Analytical Sciences). All values were normalized to empty vector (EV) to produce fold induction values.

### Quantification and Statistical Analysis

All data are expressed as the mean ± s.e.m. All statistical analyses were performed using GraphPad PRISM 10. Student’s *t*-test was used for comparison between two groups. To compare differences between groups *in vivo*, a 2-way ANOVA with Bonferroni’s multiple comparison test was performed if values were derived from a normal distribution. Where a normal distribution could not be confirmed or sample size was small, nonparametric Mann–Whitney U tests with a post hoc test were performed. A *p* value of < 0.05 was considered statistically significant. The number of sample replicates and statistical cut-offs used in the analysis of genomics data are indicated in the figure legends or the text.

