## Supplementary files for "Negative regulation of T_H_17-mediated inflammation by the nuclear receptor REV-ERBβ"

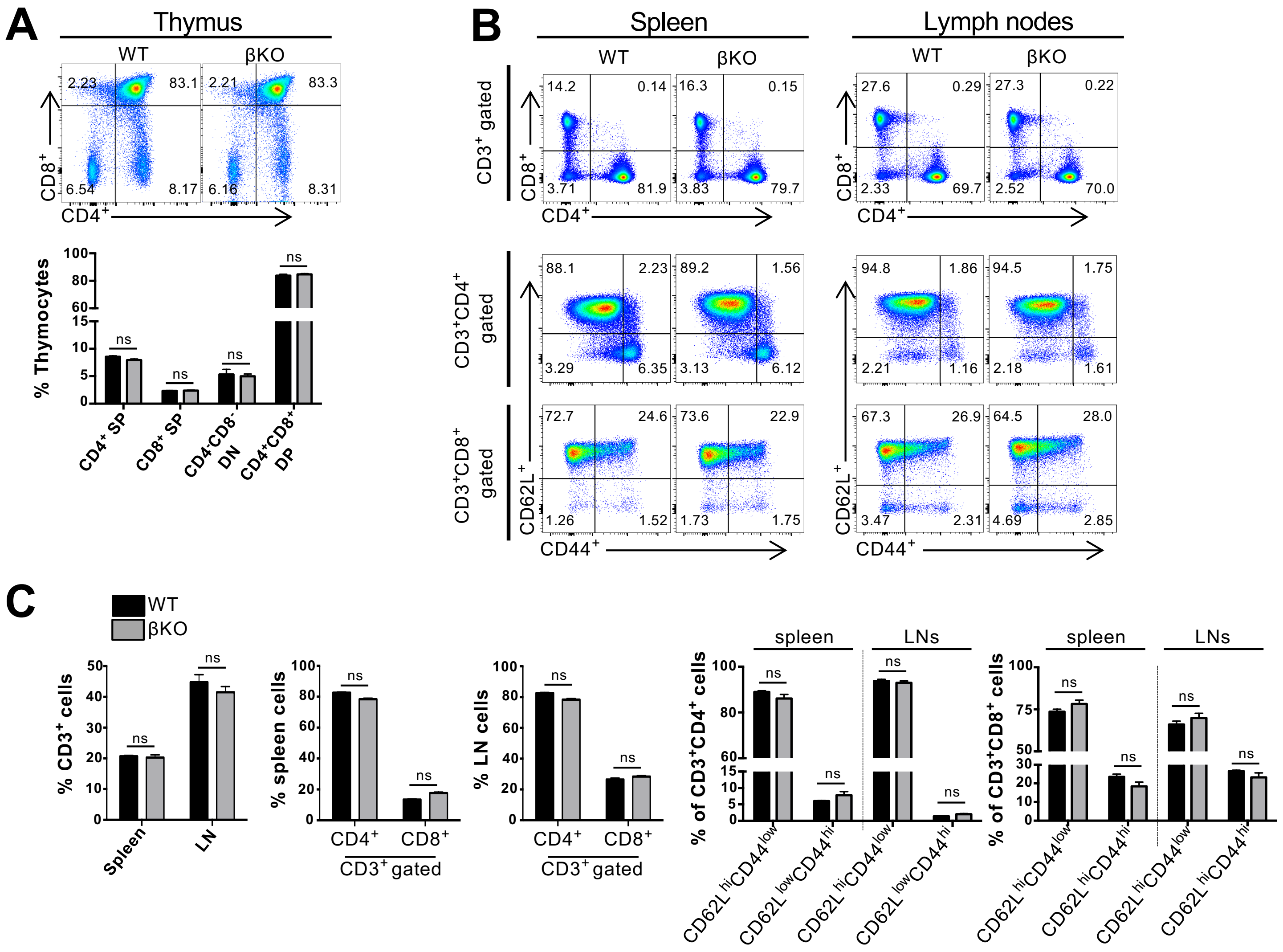


**Figure S1.** **Characterization of REV-ERBβ KO mice.** (**A**) Thymocytes from 8-week old REV-ERBβ wild-type (WT) and REV-ERBβ knock-out (βKO) mice stained with anti-CD4 and anti-CD8 antibodies and analyzed using flow cytometry. Graph (below) depicts the frequency of thymocytes per subset. (SP – single positive; DP – double positive; DN – double negative) (**B**) (*top panels*) FACS analysis of $CD3^{+}CD4^{+}$ or $CD3^{+}CD8^{+}$ T cells in the spleen and lymph nodes of WT or βKO mice. (*bottom panels*) FACS analysis of naïve ($CD62L^{high}CD44^{low}$) and effector T cell ($CD62L^{low}CD44^{high}$) populations from the spleen and lymph nodes of WT or βKO mice. (**C**) Graphs depicting the frequencies of T cells and their subsets in the spleens and lymph nodes of WT or βKO mice shown in Panel B. Data represent mean±s.e.m. (n=3/group). ns, not significant (*$p>0.05$*).

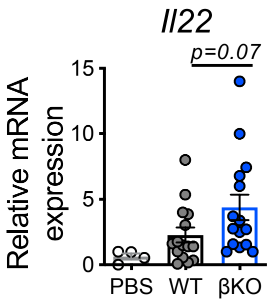


**Figure S2**. **REV-ERBβ-deficiency leads to enhanced IL-22 expression *in vivo*.** qRT-PCR analysis of *Il22* expression in the proximal colon from *Rag1*$^{-/-}$ mice receiving PBS, WT, or βKO CD4$^{+}$ T cells. *18s* was used as the internal control. Each symbol represents an individual mouse. (n=13 for WT, n=15 for βKO, and n=4 for PBS). Data represents mean±s.e.m. Student's *t*-tests were performed for statistical analysis.

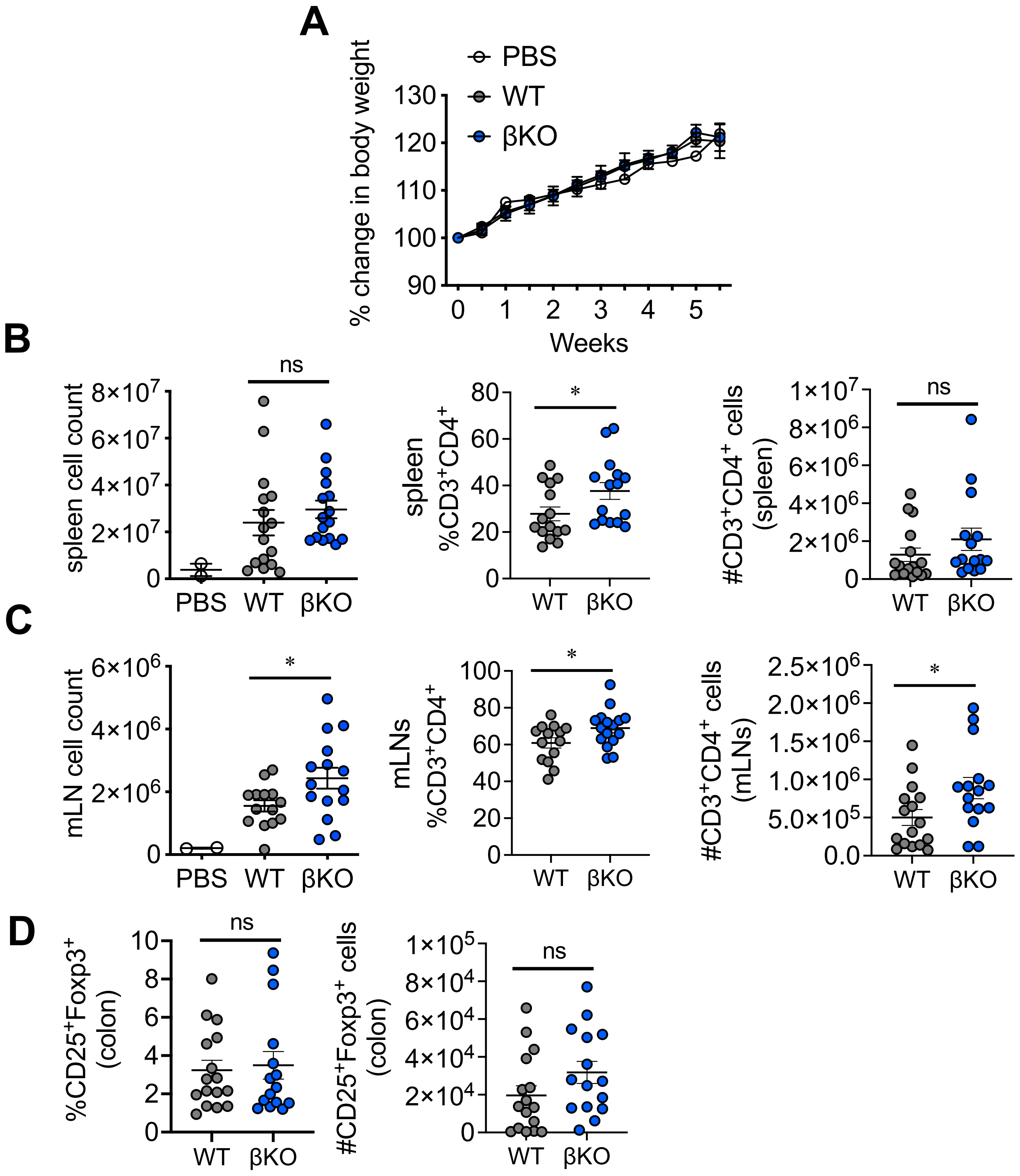


**<u>Figure S3</u>. Analysis of REV-ERBβ-deficient T cells at peak of disease. (A)** Graph depicts percent change in body weight of *Rag1*$^{-/-}$ mice receiving REV-ERBβ wild-type (WT), knock-out (βKO) T cells, or control (no cells, PBS) for the short-term (6 week) experiment. **(B)** Graphs summarizing number of cells found in the spleen (*left*), frequency, and number of CD3$^{+}$CD4$^{+}$ T cells found in the spleens (*middle, right*, respectively) of *Rag1*$^{-/-}$ mice receiving WT, βKO T cells, or PBS after 6 weeks. **(C)** Graphs summarizing number of cells found in the mLNs (*left*), frequency, and number of CD3$^{+}$CD4$^{+}$ T cells found in the mLNs (*middle, right*, respectively of *Rag1*$^{-/-}$ mice receiving WT, βKO T cells, or PBS after 6 weeks. **(D)** Graphs summarizing % and number of Foxp3$^{+}$ T regulatory cells found in the of *Rag1*$^{-/-}$ mice receiving WT and βKO T cells after 6 weeks. Each symbol represents an individual mouse. (n=14 for WT, n=15 for βKO, and n=2 for PBS). Data represents mean±s.e.m. Two-way ANOVA (body weight) and Student's *t*-tests were performed for statistical analysis. *$p<0.05$; ns, not significant.

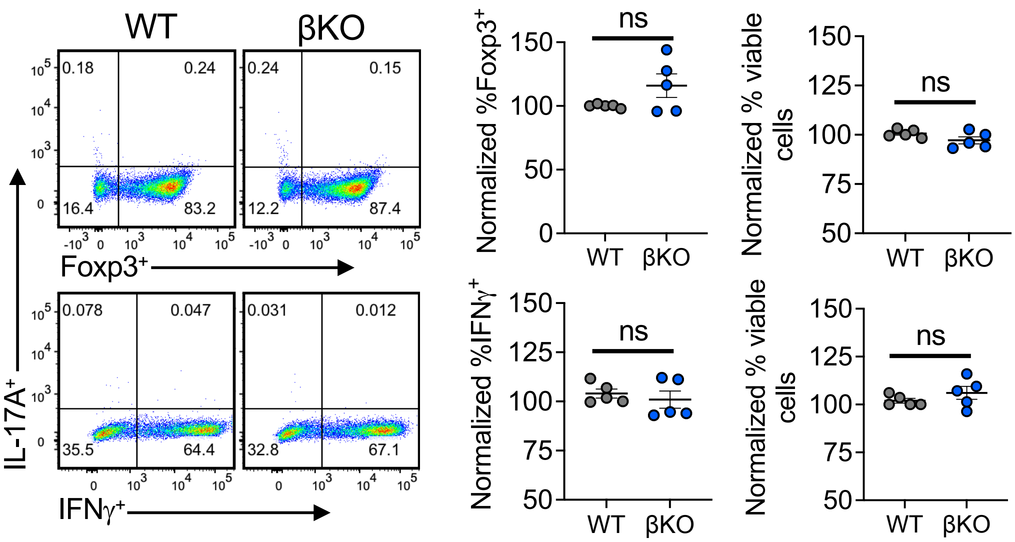


**Figure S4**. **Loss of REV-ERBβ does not affect the development of $T_H1$ and iTreg cells *in vitro*.** FACS analysis of naïve CD4⁺ T cells differentiated into $T_H1$ or iTreg cells from WT or βKO mice. FACS plots show IL-17A, IFNγ, and Foxp3 expression in the cultures. Graphs to the right show normalized IFNγ or Foxp3 expression as well normalized viability in all cell cultures (n=5). Statistics determined using Student's *t*-test. ns, not significant.

**Supplementary Table 1. qRT-PCR primers**

| Gene name | Forward primer (5' - 3') | Reverse Primer (5' - 3') |
|---|---|---|
| *b-actin* | CCACAGCTGAGAGGGAAATC | AAGGAAGGCTGGAAAAGAGC |
| *18s* | GTAACCCGTTGAACCCCATT | CCATCCAATCGGTAGTAGCG |
| *Cxcl10* | TCCTTGTCCTCCCTAGCTCA | ATAACCCCTTGGGAAGATGG |
| *Ifng* | TGGCTGTTTCTGGCTGTTACT | GCTCTGCAGGATTTTCATGTC |
| *Il6* | CATGTTCTCTGGGAAATCGTG | TCCAGTTTGGTAGCATCCATC |
| *Il17a* | CTCCAGAAGGCCCTCAGACTAC | AGCTTTCCCTCCGCATTGACACAG |
| *Il17f* | GAGGATAACACTGTGAGAGTTGAC | GAGTTCATGGTGCTGTCTTCC |
| *Il22* | TCA TCG GGG AGA AAC TGT TC | CAT GTA GGG CTG GAA CCT GT |
| *Nr1d1* | ACCTTTGAGGTGCTGATGGT | CTCGCTGAAGTCAAACATGG |
| *Nr1d2* | TGTGAAAACAGGCAAAACCA | CCCTTACAGCCTTCACAAGC |
| *Rorc* | CCGCTGAGAGGGCTTCAC | TGCAGGAGTAGGCCACATTACA |
| *Tnfa* | TCAGCCGATTTGCTATCTCAT | TGGAAGACTCCTCCCAGGTAT |
